# Dopamine Depletion Alters Macroscopic Network Dynamics in Parkinson’s Disease

**DOI:** 10.1101/382994

**Authors:** James M. Shine, Peter T. Bell, Elie Matar, Russell A. Poldrack, Simon J.G. Lewis, Glenda M. Halliday, Claire O’Callaghan

**Affiliations:** Brain and Mind Centre, The University of Sydney, Sydney, NSW, Australia; The University of Queensland, Brisbane, QLD, Australia; Department of Psychiatry and Behavioural and Clinical Neuroscience Institute, University of Cambridge, Cambridge, UK; Department of Psychology, Stanford University, Stanford, CA, USA

## Abstract

**Abstract:** Parkinson’s disease is primarily characterised by diminished dopaminergic function, however the impact of these impairments on large-scale brain dynamics remains unclear. It has been difficult to disentangle the direct effects of Parkinson’s disease from compensatory changes that reconfigure the functional signature of the whole brain network. To examine the causal role of dopamine depletion in network-level topology, we investigated time-varying network structure in 37 individuals with idiopathic Parkinson’s disease, both ‘On’ and ‘Off’ dopamine replacement therapy, along with 50 age-matched, healthy control subjects using resting-state functional MRI. By tracking dynamic network-level topology, we found that the Parkinson’s disease ‘Off’ state was associated with greater network-level integration than in the ‘On’ state. The extent of integration in the ‘Off’ state inversely correlated with motor symptom severity, suggesting that a shift toward a more integrated network topology may be a compensatory mechanism associated with preserved motor function in the dopamine depleted ‘Off’ state. Furthermore, we were able to demonstrate that measures of both cognitive and brain reserve (i.e., premorbid intelligence and whole brain grey matter volume) had a positive relationship with the relative increase in network integration observed in the dopaminergic ‘Off’ state. This suggests that each of these factors plays an important role in promoting network integration in the dopaminergic ‘Off’ state. Our findings provide a mechanistic basis for understanding the PD ‘Off’ state and provide a further conceptual link with network-level reconfiguration. Together, our results highlight the mechanisms responsible for pathological and compensatory change in Parkinson’s disease.

## Introduction

Parkinson’s disease (PD) is a common neurological disorder characterised by degeneration of the dopaminergic midbrain. This pathological insult to the brainstem results in a severe dopamine depletion throughout ascending neural pathways innervating the basal ganglia, thalamus and cortex (Braak *et al*., 2004). The impact of such extensive dopaminergic loss on brain network dynamics remains poorly understood, partly due to the fact that dopamine depletion has been linked to both pathological and compensatory changes in brain network organisation and connectivity (Bell *et al*., 2015; Bohnen and Martin, 2014; Poston *et al*., 2017).

Studies utilising resting state fMRI in PD consistently show alterations in functional connectivity that impact a diverse set of brain regions, including both cortico-cortical and cortico-subcortical architectures. In the dopaminergic ‘Off’ state, cortico-striatal hyper-connectivity is often observed, particularly in motor networks involving the subthalamic nucleus and primary motor cortex (Baudrexel *et al*., 2011; Kwak *et al*., 2012; Wu *et al*., 2010). However, alterations in inter-striatal connectivity, and across a range of cortico-striatal networks, have also been shown (Bell *et al*., 2014; Helmich *et al*., 2010; Wu *et al*., 2010). Importantly, many of these abnormalities are normalised with dopamine replacement pharmacotherapy (Kwak *et al*., 2012; Wu *et al*., 2010), suggesting that dopamine medication may play a role in both correcting pathological activity and alleviating compensatory reorganisation.

The effects of pharmacological manipulation on functional brain network architecture are often non-linear (Brezina, 2010; Marder, 2012; Tahmasian *et al*., 2015). For instance, increases in neural activity when ‘Off’ dopaminergic therapy may reflect the compensatory engagement of non-dopaminergic systems of the brainstem or network reorganisation across the cortex and subcortex. This concept is supported by a general principal of compensation observed in ageing and neurodegeneration, wherein relatively spared circuits and networks are over-engaged to support dysfunctional nodes (Grafman, 2000). In PD this effect can be observed as shifts in the topography of cortico-striatal connectivity. For example, less dopaminergically-depleted striatal zones (such as the anterior putamen) may increase their coupling with cortical sensorimotor areas to overcome relatively severe posterior striatal dopamine pathology (Hacker *et al*., 2012; Helmich *et al*., 2010). Alternatively, cortico-cerebellar connections may be increasingly engaged to offset impaired cortico-striatal function (O’Callaghan *et al*., 2016; Wu and Hallett, 2013). When these instances of hyper-connectivity are associated with preserved behavioral function, this implies an adaptive reallocation of activity in response to focal pathological changes (Hillary and Grafman, 2017). It is also possible that in some instances, functional circuit reorganisation may represent a pathological loss of network segregation or specialisation (Fornito *et al*., 2015; Hillary and Grafman, 2017). This could conceivably occur as a “knock-on” effect at the cortical network level, stemming from a fundamental loss of segregation in basal ganglia sub-circuits (Nieuwhof *et al*., 2017), which in turn would cause an increase in correlated activity in previously segregated neural populations (Bar-Gad *et al*., 2003; C. J. Wilson, 2013).

Whether the changes are compensatory or pathological, the current mechanisms supporting rearrangement of large-scale cortical patterns in the dopamine-depleted state are unclear. Ultimately, the degree of compensatory *versus* maladaptive change may be determined by the relative balance between focused increases in connectivity and a more general loss of segregation. Importantly, this concept can be examined using network analytic approaches. Using the mathematical formalism of graph theory, network communities are taken to represent densely interconnected neural elements in which local connections are highly segregated. In contrast, network hubs integrate diverse communities, enabling channels for effective information integration (Bertolero *et al*., 2017; van den Heuvel and Sporns, 2013). These organisational principles are thought to balance the specialisation of function with the integration of information (Deco *et al*., 2015; Park and Friston, 2013), and this balance gives rise to complex neural dynamics that span multiple spatiotemporal scales (Deco *et al*., 2013; Honey *et al*., 2012).

Here, we used time-resolved functional connectivity of resting state fMRI in combination with graph theoretical analyses to determine the balance between integration and segregation in the face of dopaminergic depletion in Parkinson’s disease. To date, studies of network topology in neurological disease have largely focused on structural brain networks and time-averaged resting-state functional networks, which represent an inherently ‘static’ snapshot of brain architecture (Breakspear, 2017). However, recent advances in the statistical analysis of time-varying resting-state functional MRI data have demonstrated that functional brain organisation is dynamic over the course of seconds to minutes (Betzel et al., 2016; Shine et al., 2016; Zalesky et al., 2014) and that fluctuating network dynamics are crucial for normal cognitive (Hearne *et al*., 2017; Shine *et al*., 2016) and motor (Bassett *et al*., 2011) function. Examination of time varying functional network architecture provides an opportunity to explore the balance between segregated and integrated neural dynamics in both health and disease.

In determining the impact of dopamine depletion on dynamic network architecture in PD, we further aimed to establish whether certain functional patterns in the ‘Off’ state may be linked to a compensatory mechanism. Defining compensatory activity in neurodegeneration is a nontrivial problem (Gregory *et al*., 2017). Brain and cognitive reserve, respectively, refer to aspects of structural integrity that support increased functional resilience, and the preservation of function in the face of underlying degeneration (Fratiglioni and Wang, 2007). Importantly, these concepts can be operationalised using grey matter integrity (as a surrogate of brain reserve) and educational level or general intelligence quotient (as a surrogate of pre-morbid cognitive reserve) (Stern, 2017). These metrics can then be compared to network-level topological changes to provide an estimate of the extent to which brain organisation is related to functional and structural resilience.

To examine the dynamic network architecture of the resting brain in the ‘Off’ compared with the ‘On’ dopaminergic state, we related network topology to motor function in the ‘Off’ state, and also to measures of cognitive and brain reserve. We hypothesised that removal of dopaminergic medications would lead to a relatively integrated network topology, which should also relate to a preservation of motor function in the ‘Off’ state. In addition, any compensatory pattern should be observed in the positive relationship between network topology and both cognitive and brain reserve.

## Materials and Methods

### Participants

37 patients were recruited from the Parkinson’s Disease Research Clinic at the Brain and Mind Centre, University of Sydney, Australia. All patients satisfied the United Kingdom Parkinson’s Disease Society Brain Bank criteria and were not demented (Martinez-Martin *et al*., 2011). Patients were assessed on the Hoehn and Yahr Scale and the motor section of the unified Parkinson’s disease rating scale (UPDRS-III) in the dopaminergic ‘Off’ state. The Mini-mental state examination (MMSE) was administered as a measure of general cognition.

Participants with Parkinson’s disease were assessed on two occasions: ‘On’ their regular dopaminergic medications and ‘Off’ following overnight withdrawal (i.e. 12-18 hours) of dopaminergic medications (5.2 ± 1.4 weeks between sessions). Dopaminergic dose equivalence (DDE) scores were calculated for each patient. Specifically, 10 patients were on L-dopa monotherapy; 9 were on L-dopa plus a dopaminergic agonist; a further 8 were on L-dopa plus adjuvant therapy (rasagaline, entacapone or a monoamine oxidase inhibitor); 7 were on a combination of L-dopa, dopaminergic agonist and adjuvant therapy; one patient was on dopaminergic agonist monotherapy, and two were on an agonist plus adjuvant therapy. No participant was taking any psychoactive medications.

50 healthy controls were recruited to participate in the study. Control participants were screened for a history of neurological or psychiatric disorders, and no controls were using psychoactive medications. Patients with PD and healthy controls were matched for age and education. The study was approved by the local Ethics Committees and all participants provided informed consent in accordance with the Declaration of Helsinki. See Table 1 for demographic details and clinical characteristics.

**Table 1.** Mean (standard deviation) for demographics and patient clinical characteristics.

|  | Control | PD |
| --- | --- | --- |
| N | 50 | 37 |
| Sex (M:F) | 14:36 | 29:8 |
| Age | 65.82 (7.8) | 65.05 (7.2) |
| Education | 13.51 (2.8) | 13.50 (3.0) |
| MMSE | 29.00 (1.2) | 28.59 (2.2) |
| BDI-II | 9.41 (7.3) | 10.67 (8.6) |
| Duration (yrs diagnosed) | - | 6.41 (4.2) |
| DDE (mg/day) | - | 808.54 (503.0) |
| Hoehn & Yahr stage | - | 2.14 (0.8) |
| UPDRS III, 'Off' | - | 32.00 (16.3) |
| NART | - | 111.08 (10.3) |

MMSE = Mini-Mental State Examination; DDE = Dopaminergic dose equivalence; UPDRS III = Motor section of the unified Parkinson's disease rating scale; BDI-II = Beck Depression Inventory-
II. There were no significant differences in age, education, MMSE or BDI-II between the two groups, though there were differences in gender ( $p < 0.05$ ).

### Behavioural and neuropsychological assessment

Mood was assessed via a self-report questionnaire, the Beck Depression Inventory-II (BDI-II; Beck *et al*., 1996). Patients were also administered the National Adult Reading Test (NART (Bright *et al*., 2016), and their predicted pre-morbid full scale IQ was calculated. The NART is an established measure of premorbid intelligence and serves as a surrogate of cognitive reserve, with the benefit of offering greater variance than years of education in a homogenous sample (Stern *et al*., 2003). These measures were assessed in the dopaminergic ‘On’ state. Results from these measures are also shown in Table 1.

### Imaging acquisition

Imaging was conducted on a General Electric 3 Tesla MRI (General Electric, Milwaukee, USA). Whole-brain three dimensional T1-weighted sequences were acquired as follows: coronal orientation, matrix 256 x 256, 200 slices, 1 x 1 mm^2^ in-plane resolution, slice thickness 1 mm, TE/TR = 2.6/5.8 ms. T2^*^-weighted echo planar functional images were acquired in interleaved order with repetition time (TR) = 3 s, echo time (TE) = 32 ms, flip angle 90°, 32 axial slices covering the whole brain, field of view (FOV) = 220 mm, interslice gap = 0.4 mm, and raw voxel size = 3.9 mm by 3.9 mm by 4 mm thick. Each resting state scan lasted 7 min (140 TRs). During the resting-state scan, patients were instructed to lie awake with their eyes closed and to let their minds wander freely.

### Resting state fMRI data

Preprocessing and analyses of resting state data were conducted using SPM12 (http://www.fil.ion.ucl.ac.uk/spm/software/). Scans were first slice-time corrected to the median slice in each TR, then realigned to create a mean realigned image, with measures of 6 degrees of rigid head movements calculated for later use in the correction of minor head movements. For quality assurance, each trial was analyzed using ArtRepair (Mazaika *et al*., 2009) and trials with a large amount of global drift or scan-to-scan head movements greater than 1 mm were corrected using interpolation. None of the subjects included in this study demonstrated scan-to-scan head movements >3mm (less than one voxel breadth). Images were normalized to the Echo Planar Image template, resampled to 3mm isotropic voxels and then subsequently smoothed using a 4mm full-width half-maximum isotropic Gaussian kernel.

Temporal artifacts were identified in each dataset by calculating framewise displacement (FD) from the derivatives of the six rigid-body realignment parameters estimated during standard volume realignment (Power *et al*., 2014), as well as the root mean square change in BOLD signal from volume to volume (DVARS). Frames associated with FD > 0.25mm or DVARS > 2.5% were identified, however as no participants were identified with greater than 10% of the resting time points exceeding these values, no sessions were excluded from further analysis. However, to ensure that neither head motion or the global signal were responsible for any group effects, we re-ran the analysis after: i) scrubbing data with FD > 0.25mm or DVARS > 2.5%; or ii) global signal regression. Both analyses revealed similar group-level effects.

Following artifact detection, nuisance covariates associated with the 12 linear head movement parameters (and their temporal derivatives), FD, DVARS, and anatomical masks from the CSF and deep cerebral WM were regressed from the data using the CompCor strategy (Behzadi *et al*., 2007). In keeping with previous time-resolved connectivity experiments (Bassett *et al*., 2015), a temporal band pass filter (0.071 < f < 0.125 Hz) was applied to the data. Finally, given the importance of head motion in functional connectivity analyses, we compared mean framewise displacement (Power *et al*., 2014) across the entire resting state session across the three groups (controls, PD ‘On’ and PD ‘Off’).

### Brain parcellation

Following pre-processing, the mean time series was extracted from 347 predefined parcels. To ensure whole-brain coverage, we extracted: 333 cortical parcels (161 and 162 regions from the left and right hemispheres, respectively) using the Gordon atlas (Gordon *et al*., 2014) and 14 subcortical regions from Harvard-Oxford subcortical atlas (bilateral thalamus, caudate, putamen, ventral striatum, globus pallidus, amygdala and hippocampus; http://fsl.fmrib.ox.ac.uk/) for each participant in the study.

### Time-resolved functional connectivity

To estimate functional connectivity between the 347 parcels, we used the Multiplication of Temporal Derivatives; MTD metric (Shine *et al*., 2015). The MTD is computed by calculating the point-wise product of temporal derivative of pairwise time series (Equation 1). The MTD is averaged by calculating the mean value over a temporal window, *w*. Time-resolved functional connectivity was calculated between all 347 brain regions using the MTD within a sliding temporal window of 15 time points (~33 seconds), which allowed for estimates of signals amplified at approximately 0.1 Hz (Shine *et al*., 2015). Individual functional connectivity matrices were then calculated within each temporal window, thus generating a weighted 3-dimensional adjacency matrix (region × region × time) for each participant.

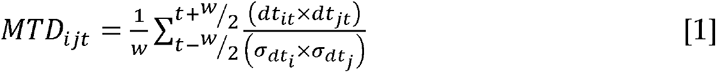

**Equation 1** – Multiplication of Temporal Derivatives, where for each time point, *t*, the MTD for the pairwise interaction between region *i* and *j* is defined according to equation *1*, where *dt* is the first temporal derivative of the *i^th^* or *j^th^* time series at time *t*, σ is the standard deviation of the temporal derivative time series for region *i* or *j* and *w* is the window length of the simple moving average. This equation can then be calculated over the course of a time series to obtain an estimate of time-resolved connectivity between pairs of regions.

### Time- resolved community structure

The Louvain modularity algorithm was applied to the functional connectivity time series using the Brain Connectivity Toolbox (Rubinov and Sporns, 2010). The Louvain algorithm iteratively maximizes the modularity statistic, Q, for different community assignments until the maximum possible score of *Q* has been obtained (Equation 2). The modularity estimate for a given adjacency matrix quantifies the extent to which the network may be subdivided into communities with stronger within-module than between-module connections. Using this technique, time-averaged and time-resolved community structure was calculated for each participant.

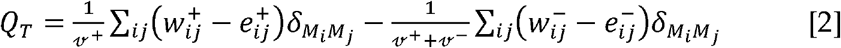

**Equation 2** – Louvain modularity algorithm, where *v* is the total weight of the network (sum of all negative and positive connections), *w_ij_* is the weighted and signed connection between regions *i* and *j*, *e_ij_* is the strength of a connection divided by the total weight of the network, and *δ_MiMj_*, is set to 1 when regions are in the same community and 0 otherwise. ‘+’ and superscripts denote all positive and negative connections, respectively.

For each temporal window, regional community assignment was assessed 500 times and a consensus partition was identified using a fine-tuning algorithm from the Brain Connectivity Toolbox. This then afforded an estimate of both the time resolved modularity (*Q_T_*) and cluster assignment (Ci_T_) within each temporal window for each participant in the study. To define an appropriate value for the γ parameter, we iterated the Louvain algorithm across a range of values (0.5-2.5 in steps of 0.1) for 100 iterations of a single subject’s (sub1) time-averaged connectivity matrix and then estimated the similarity of the resultant partitions using mutual information. A γ parameter of 1.1 provided the most robust estimates of topology across these iterations (quantified by the minimum standard deviation across 100 iterations of the Louvain algorithm).

### Cartographic profiling

Based on time-resolved community assignments, we estimated within-module connectivity by calculating the time-resolved module-degree Z-score (W*_T_*; within module strength) for each parcel (Equation 3) (Guimerà and Nunes Amaral, 2005).

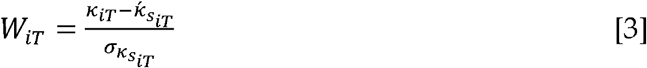

**Equation 3** – Module degree Z-score, *W_iT_* where κ_i_T is the strength of the connections of region *i* to other regions in its module s_i_ at time *T*, 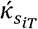 is the average of κ over all the regions in si at time *T*, and 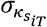 is the standard deviation of κ in si at time *T*.

To calculate between-module connectivity (B*_T_*), we used the participation coefficient, B*_T_*, quantifying the extent to which a region connects across all modules (i.e. between-module strength; equation 4).

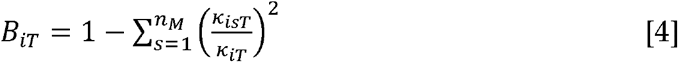

**Equation 4** – Participation coefficient B*iT*, where κ_is_T is the strength of the positive connections of region *i* to regions in module s at time *T*, and κ_iT_ is the sum of strengths of all positive connections of region *i* at time *T*. The participation coefficient of a region is therefore close to 1 if its connections are uniformly distributed among all the modules and 0 if all of its links are within its own module.

To track fluctuations in cartography over time, for each temporal window, we computed a joint histogram of within- and between-module connectivity measures, referred to here as a “cartographic profile” (Figure 1b). Code for this analysis is freely available at https://github.com/macshine/integration/.

**Figure 1.**
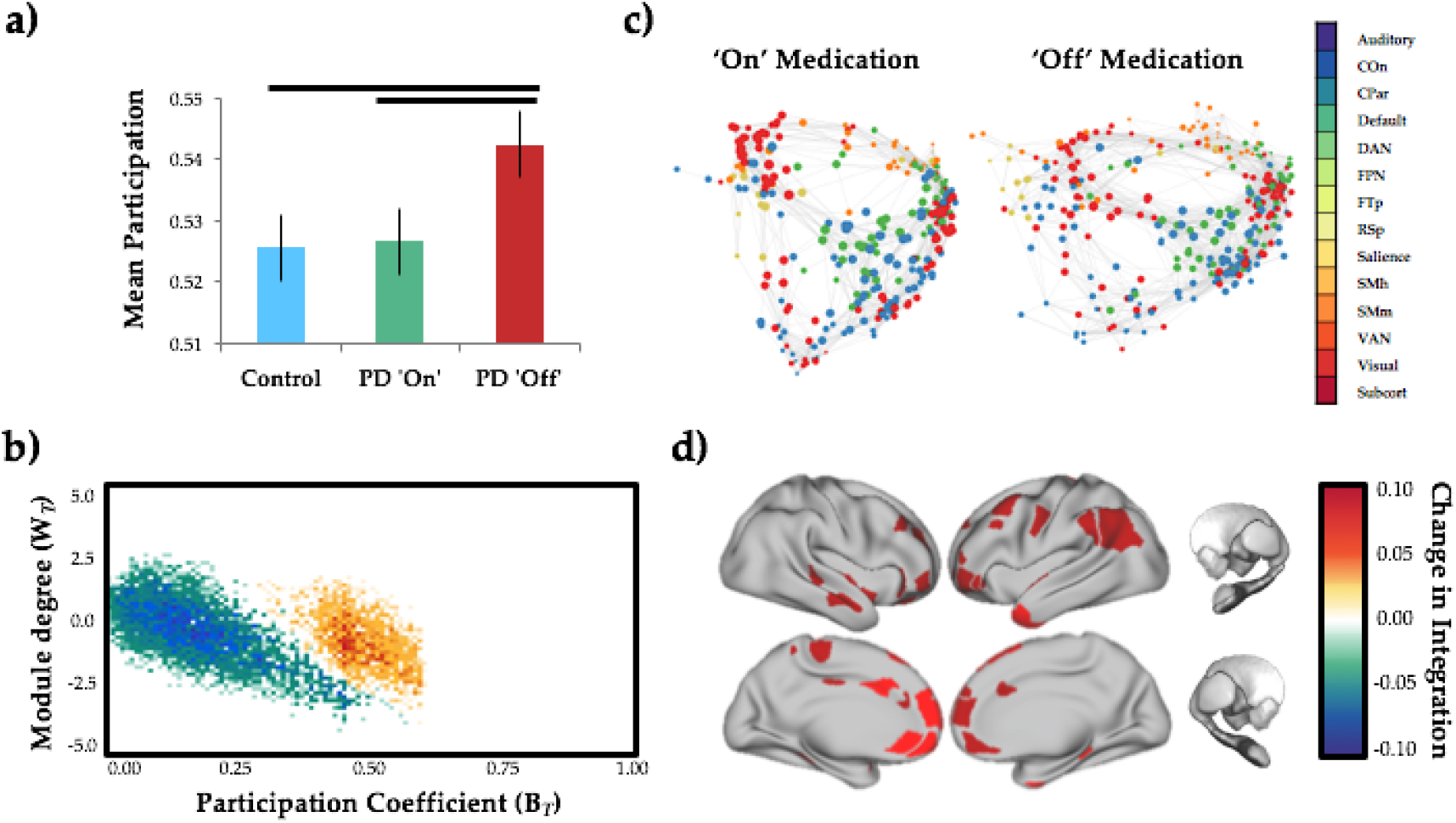
Network topology as a function of dopaminergic state.

a) global mean participation coefficient (B*_T_*) in controls (blue), PD ‘On’ (teal) and PD ‘Off’ (red) – p < 0.001; b) cartographic profile comparing PD ‘Off’ > PD ‘On’ – subjects were more integrated (i.e. rightward shift on the B*_T_* axis) in the ‘Off’ compared to the ‘On’ state; c) force-directed plots comparing PD ‘On’ and ‘Off’ dopaminergic medication – edges represent top 1% of connections in time-averaged connectivity matrix and colors of nodes reflect pre-defined network identity of each region; d) surface plot of regions with significantly increased participation (B*_T_*) during ‘Off’ state.

### Regional Flexibility

The flexibility of each brain parcel was calculated by the percentage of temporal windows in which an individual region ‘switched’ between modules, normalized to the total number of modules in the data (as estimated in the previous step). Code was obtained directly from the original author (http://www.danisbassett.com/resources/). As the modular assignment was essentially arbitrary within each unique temporal window, we used a version of the Hungarian algorithm to assign regions to modules with consistent values over time.

### Grey matter extraction

Grey matter extraction was performed using the FMRIB software library package FSL (http://www.fmrib.ox.ac.uk/fsl/). Scans were skull-stripped using the BET algorithm in FSL (Smith, 2002) and tissue segmentation was completed using FMRIB’s Automatic Segmentation Tool (FAST v4.0) (Zhang *et al*., 2001). A study-specific grey matter template was created using the maximum equal number of scans from both groups (37 from each) and registered to the Montreal Neurological Institute Standard space (MNI 152) using a non-linear b-spline representation of the registration warp field. Grey matter partial volume maps were non-linearly registered to the study template and modulated by dividing by the Jacobian of the warp field, to correct for any contraction/enlargement caused by the non-linear component of the transformation; this step corrects for total intracranial volume (ICV) so that it does not need to be included as a confounding covariate (Good *et al*., 2002). After normalisation and modulation, the grey matter maps were smoothed with an isotropic Gaussian kernel with a sigma of 2 mm.

Whole brain grey matter volume (mm^3^) was then extracted for each participant. The total volume of non-zero voxels was extracted from the grey matter mask automatically generated from FAST. Using the smoothed and registered images, the mean proportion of grey matter per voxel from non-zero voxels was extracted for each subject using fslstats. Multiplying the volume within the mask by each subjects’ mean grey matter proportion inside the mask gave a measure of total grey matter volume for each person. We used whole brain grey matter volume (corrected for total ICV), a specific indicator of grey matter structural integrity, as our measure of brain reserve. For completeness, we also calculated total ICV, as it is a commonly used proxy for brain reserve. Total ICV was calculated for each individual by summing the segmented grey matter, white matter and cerebrospinal fluid volumes obtained from the FAST procedure. We re-ran our analysis using total ICV as a measure of brain reserve to confirm that similar results were obtained using both total grey matter volume and total ICV as indicators of brain reserve.

### Statistical analyses

To determine whether there were any abnormalities in functional network topology between groups, the mean cartographic profile for each PD patient was compared between medication states (paired-sample t-test for each bin of the cartographic profile; FDR q ≤ 0.05) and between groups (independent-samples t-test for each bin; FDR q ≤ 0.05). Regional W_T_ and B_T_ scores were also compared across groups using independent-samples t-tests.

To determine the clinical relevance of functional network reconfiguration, we measured the correlation of the difference in the cartographic profile between the ‘Off’ and ‘On’ state with the severity of motor impairments in the ‘Off’ state (measured using UPDRS III) using a Spearman’s rho correlation (due to the non-parametric nature of the data).

To determine whether network level integration related to brain reserve in the individuals with PD, we fit a general linear model which fit grey matter volume, predicted full scale IQ (as estimated using normalised National Adult Reading Test [NART] scores) and the interaction between these two measures of reserve (i.e. GM*NART) to the amount of integration present in the ‘Off’ vs ‘On’ state, while co-varying for age. Separate analyses were conducted at the global (i.e. cartographic profile) and regional (i.e. parcel-wise) level.

## Results

### Head motion

There were no significant differences in head movement between healthy controls and individuals with PD in either medication state (mean framewise displacement: controls 4.9×10^-4^ ± 3.3×10^-4^, PD ‘On’: 5.7×10^-4^ ± 5.3×10^-4^; PD ‘Off’: 5.8×10^-4^ ± 5.0×10^-4^; all p > 0.200) or between dopaminergic states in the PD group (p > 0.200).

### Relationship between network topology and dopaminergic state

In the dopaminergic ‘Off’ state, individuals with PD demonstrated a more integrated functional network topology than those ‘On’ medication or healthy controls (Figure 1a,b; p < 0.001). The magnitude of regional between-module integration was significantly higher in the ‘Off’ state (relative to the ‘On’ state) across medial and lateral frontoparietal cortical regions (Figure 1b). These changes were diffusely mediated across multiple sub-systems, including frontoparietal, cingulo-opercular, salience and dorsal attention networks (Figure 1c,d).

### Relationship between network topology and motor symptom severity

There was a significant inverse correlation between network-level integration and UPDRS III scores as measured in the ‘Off’ state (Figure 2a) that was maximal in right dorsolateral prefrontal cortex, bilateral dorsal anterior cingulate, bilateral retrosplenial cortex and sensorimotor cortex (Figure 2b). This is consistent with greater integration in those regions being associated with less severe motor impairment on UPDRS motor scale.

**Figure 2.**
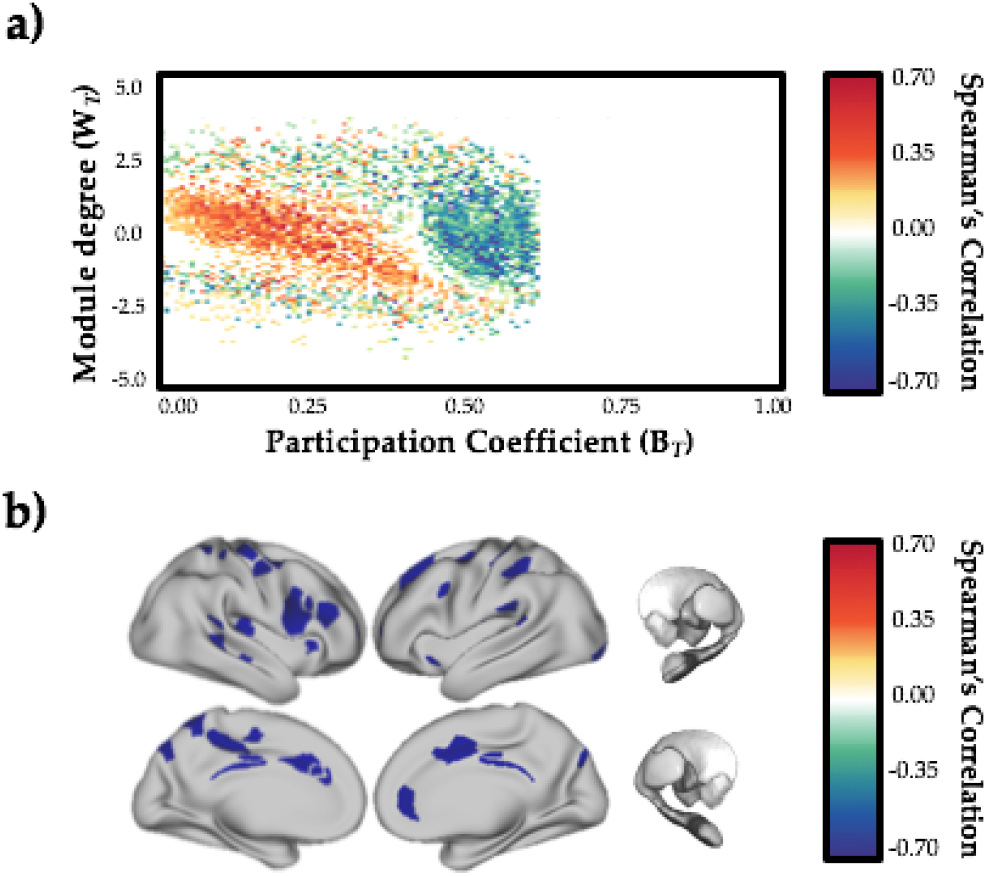
Relationship between network topology and motor severity.

a) inverse relationship between cartographic profile (PD ‘Off’ > PD ‘On’) and UPDRS III (motor) severity (estimated in the dopaminergic ‘Off’ state) – greater integration (i.e. rightward shift on the B*_T_* axis) was inversely correlated with motor severity; b) parcels with significant inverse correlation between B*_T_* (‘Off’ > ‘On’) and UPDRS III – FDR q ≤ 0.05.

### Effects of dopaminergic state on regional flexibility

Regions were more likely to switch their modular allegiance more frequently in the dopaminergic ‘Off’ state as compared to the ‘On’ state. We observed a distributed set of insular, frontal and parietal regions that demonstrated an increase in topological flexibility and/or decreased modular stability in the resting state following dopaminergic withdrawal (Figure 3). We did not observe a relationship between flexibility and motor severity (all p > 0.05).

**Figure 3.**
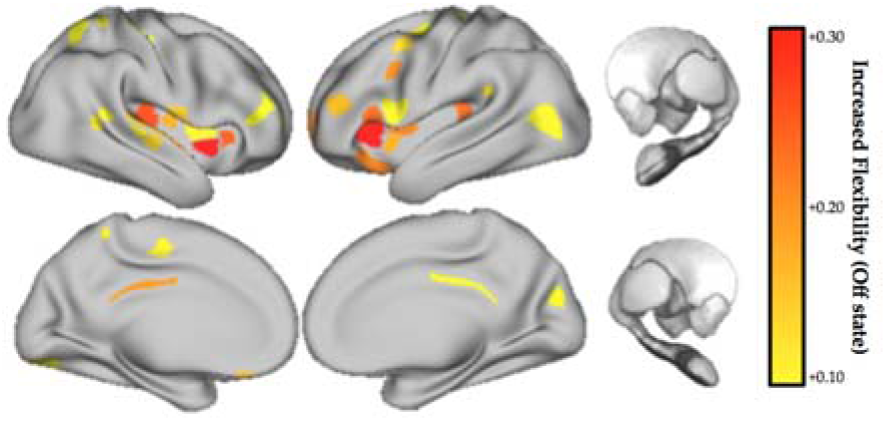
Topological flexibility as a function of dopaminergic state.

regions with increased topological flexibility (increased frequency of modular switching) in the ‘Off’ > ‘On’ dopaminergic state – FDR q ≤ 0.05. No regions showed a significant decrease in flexibility in the ‘Off’ state.

### Relationship between network topology and brain reserve

Using a linear mixed effects model specifying GM (Figure S1a), NART (Figure S1b) and GM*NART (Figure 4a) adjusting for age and gender as fixed covariates, we observed a positive relationship (FDR q ≤ 0.05) between network topology and the interaction between grey matter volume and NART-predicted IQ scores) that was maximal in frontal cortex, insula, thalamus and amygdala (Figure 4b; FDR q ≤ 0.05).

**Figure 4.**
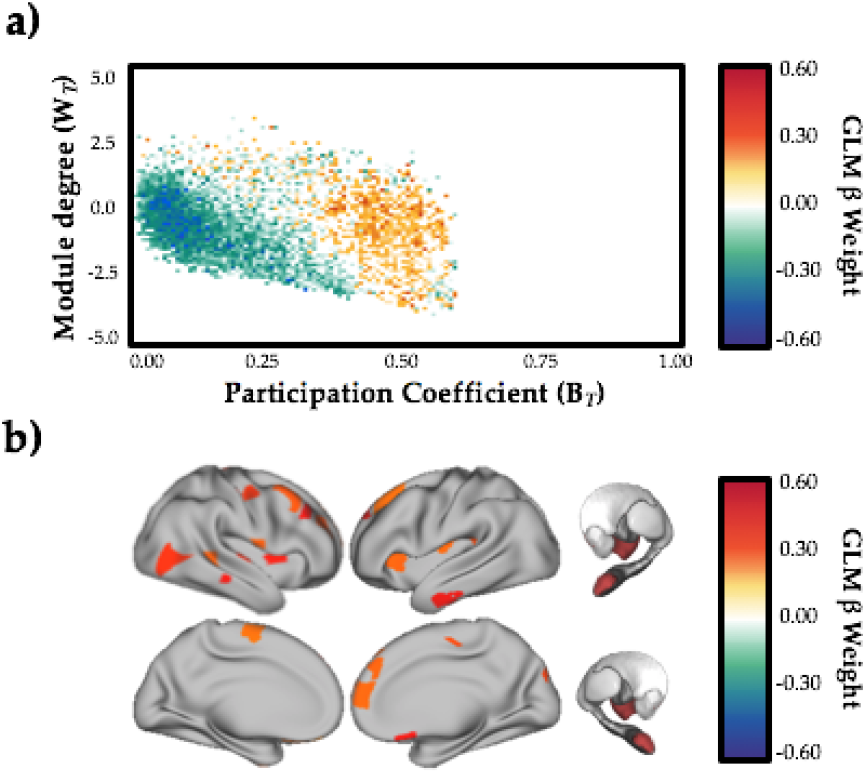
Relationship between network topology and neurocognitive reserve.

a) relationship between cartographic profile (PD ‘Off’ > PD ‘On’) and interaction between grey matter volume and education level (NART) – FDR q ≤ 0.05 – subjects with greater NART and GM scores were more integrated (i.e. rightward shift on the B*_T_* axis) in the ‘Off’ compared to the ‘On’ state; b) regions with significant relationship between B*_T_* (‘Off’ > ‘On’) and the interactions between brain (mean grey matter) and cognitive (NART) reserve, estimated using a linear mixed effects model – FDR q ≤ 0.05.

## Discussion

In this study, we demonstrated causal evidence for large-scale network reconfiguration in the ‘Off’ state in individuals with idiopathic Parkinson’s disease as compared to the dopaminergic ‘On’ state, consistent with an increase in topological integration (Figure 1) and flexibility Figure 3). Within this general shift toward a more integrated state, a distributed set of regions were inversely correlated with motor symptom severity (Figure 2), suggesting that increased integration may provide compensatory processes that offset clinical motor severity. Furthermore, we showed an association between the magnitude of integration in the ‘Off’ state and measures of grey matter volume and premorbid intelligence. This suggests that a topological shift in response to dopamine depletion is related to neurocognitive reserve (Figure 4). Together these results show that the effect of dopamine depletion in PD results in a global shift toward integration, and, that this increased integration may serve some compensatory function, the extent of which may be determined by underlying cognitive and brain reserve.

Withdrawal from dopamine replacement therapy altered network topology in the medial frontal, lateral parietal and anterior temporal cortices (Figure 1d). Importantly, these regions also exhibited an increase in topological flexibility in the ‘Off’ state, suggesting that they were not effectively “locked” into an integrated state, a result that may have argued against a possible compensatory role for increased integration in ‘Off’ state. Similar regions were inversely correlated with ‘Off’ state motor symptom severity (Figure 2b), suggesting that regional and network-level integration may help maintain motor function in the face of dopamine depletion.

The possibility that increased topological integration in the face of dopamine depletion may be associated with a compensatory function supports and extends a growing literature that highlights the importance of network level hyperconnectivity as an adaptive response to local pathological change in neurodegenerative disorders (Gregory *et al*., 2018; Hillary and Grafman, 2017; O’Callaghan *et al*., 2016). In PD, this response has previously been observed and interpreted based on static measures of resting state fMRI (Helmich *et al*., 2010; O’Callaghan *et al*., 2016; Wu *et al*., 2010). Here, we provide a description of the underlying dynamic processes that might support these enhanced activations.

Prior work has highlighted a link between increased resting state functional connectivity and markers of cognitive reserve (e.g., greater years of education) in diverse cohorts, including healthy ageing, and those with mild cognitive impairment and Alzheimer’s disease (Arenaza-Urquijo *et al*., 2013; Franzmeier *et al*., 2017; 2018). However, increased functional connectivity does not necessarily lend itself to a specific mechanistic interpretation *per se*. Using the mathematical formalism of graph theory, our results identify a relationship between premorbid intelligence and the capacity to promote functional integration, suggesting a possible dynamic mechanism that underpins the role of cognitive reserve in compensation.

The use of overall brain volume as a measure of brain reserve in our study is somewhat underspecified. Whole-brain grey matter volume incorporates a host of factors, including neuronal count, neuronal integrity and synaptic density, which jointly determine the brain’s ability to engage compensatory activity. Despite this caveat, the structural integrity of nodes (and hence, the grey matter volume) is proposed to mediate network controllability, and therefore may explain the role of brain reserve in supporting resilience of large-scale networks in ageing and neurodegeneration (Medaglia *et al*., 2017). Such nodes may indeed mediate the overall flexibility of brain networks, and allow for transitions between segregated and integrated states (Pasqualetti *et al*., 2014). Here, we identified a relationship between brain volume and the capacity to move toward a more integrated state. This result is consistent with the proposed hypothesis that brain volume may serve as a proxy for network controllability, as it captures within it a measure of the structural integrity of nodes involved in network control (Medaglia *et al*., 2017).

The prospect of compensatory network-level integration in the dopamine-depleted state raises the question of the potential mechanism for this effect. One plausible hypothesis is the relative integrity of other neuromodulatory neurotransmitter systems that contribute to global brain network dynamics (Brezina, 2010). Aside from the widespread dopaminergic loss that characterises PD, the disease is also associated with neuropathological alterations within the brainstem nuclei that supply the brain with noradrenaline (Rye and DeLong, 2003), acetylcholine (Müller and Bohnen, 2013) and serotonin (Politis and Niccolini, 2015). In the ‘Off’ state, compensatory drive may be determined by the degree of relative preservation in these nuclei and the ascending projections throughout the brain.

In the context of promoting network level integration, in healthy individuals a link has been observed between the ascending noradrenergic neuromodulatory system and global functional integration (Shine *et al*., 2016; Shine, Aburn, *et al*., 2018; Shine, van den Brink, *et al*., 2018), suggesting effective functioning of this system is crucial for modulating the gain and responsiveness of ongoing neuronal processing (Shine, Aburn, *et al*., 2018). In addition, it has been proposed that activation of the locus coeruleus noradrenergic system across the lifespan is a crucial determinant of later-life cognitive reserve (R. S. Wilson *et al*., 2013), potentially through brain derived neurotrophic factor-mediated neuroplasticity (Mather and Harley, 2016; Robertson, 2013). It follows that one possible mechanism supporting compensatory increases in integration in the dopaminergic ‘Off’ state may reflect a long-term compensatory strategy, mediated at least partially by the noradrenergic locus coeruleus. The implication is that as system begins to fail, as is the case when locus coeruleus develops high levels of alpha-synuclein (Surmeier *et al*., 2017), the compensatory reserve is lost and a failure to effectively integrate the brain unmasks the clinical severity of symptoms of Parkinson’s disease.

In addition to noradrenergic function, the multi-scale nature of the brain’s neuromodulatory network (Brezina, 2010) means it is likely that other neurotransmitter systems play a crucial role in mediating adaptive brain dynamics in the face of dopaminergic cell loss. For instance, there is a well-demonstrated loss of cholinergic cell bodies in the basal nucleus in Parkinson’s disease (Müller and Bohnen, 2013). Given the recent links between the global brain signal and ascending cholinergic activity (Turchi *et al*., 2018), it is also plausible that impairments in the cholinergic system could adversely affect the topological signature of the network, or that the relative preservation of the cholinergic system might contribute to compensatory neural dynamics. The presence of serotonergic deficits (Politis and Niccolini, 2015) further points to a complex, multi-system pathological mechanism for compensation and impairment in Parkinson’s disease.

In summary, we used a combination of time-resolved resting fMRI, graph theoretical analysis and the manipulation of dopaminergic therapy in individuals with idiopathic Parkinson’s disease to provide evidence for alterations in network topology that related to motor severity. These topological signatures demonstrated a relationship with both brain and cognitive reserve, suggesting a possible compensatory role, which may be mediated by the relative integrity of other neuromodulatory systems. Future work that disambiguates the causal relationships between neuromodulatory systems and large scale network dynamics in PD, perhaps as a function of differing disease stage, will help to better clarify this and potentially uncover new avenues for pharmacological treatments.

## Supplementary Figures

**Figure S1.**
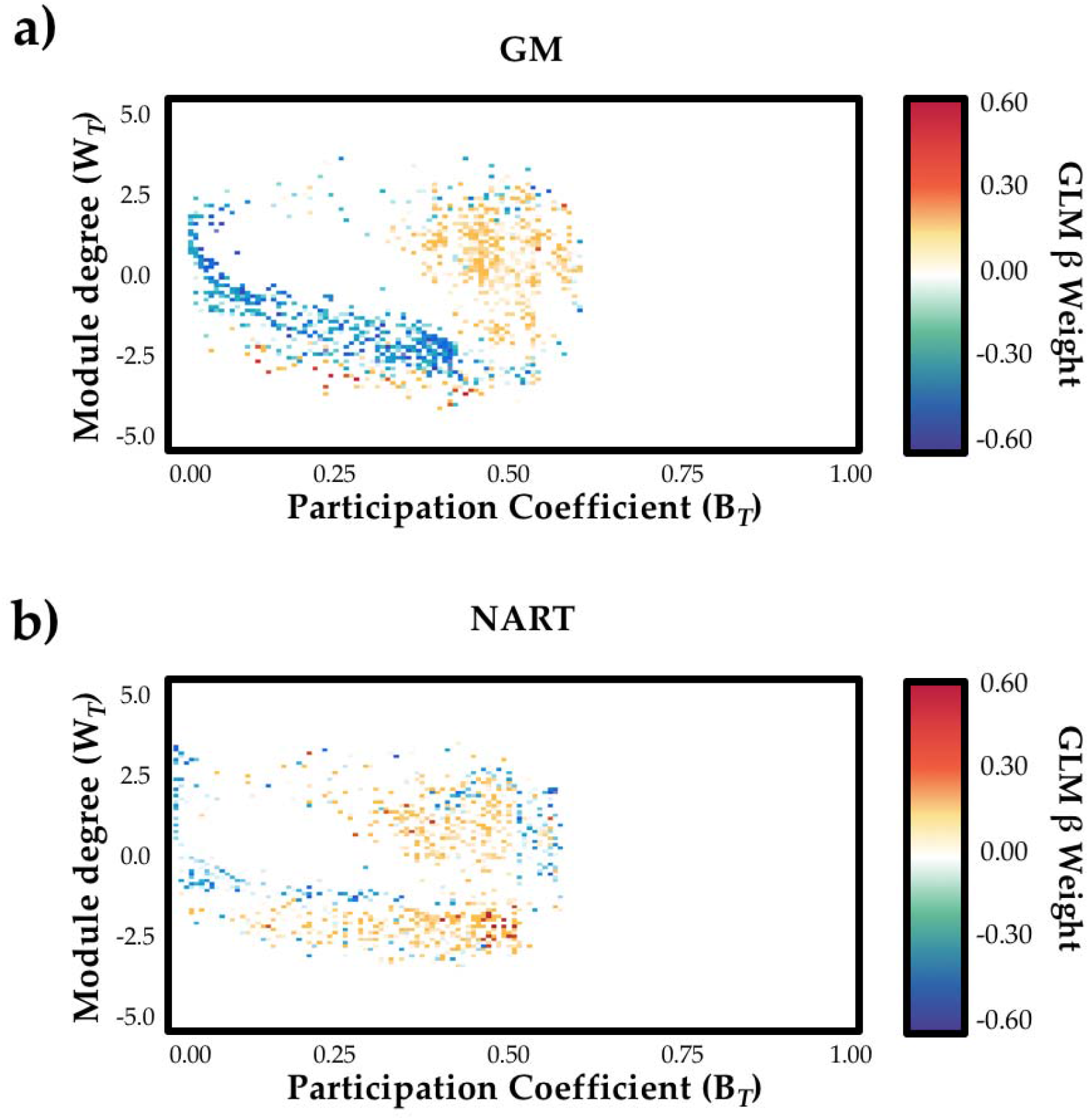
Relationship between network topology and neurocognitive reserve.

a) relationship between cartographic profile (PD ‘Off’ > PD ‘On’) and education level (NART); b) relationship between cartographic profile (PD ‘Off’ > PD ‘On’) and grey matter volume (p ≤ 0.05; not corrected for multiple comparisons).

## Acknowledgments

JMS is supported by a National Health and Medical Research Council CJ Martin Fellowship (#1072403).

EM is supported by an NHMRC Postgraduate scholarship and the Australian and New Zealand Association of Neurologists Gwen James Dementia Fellowship.

SJGL is supported by an NHMRC-ARC Dementia Fellowship (#1110414)

This work was supported by funding to Forefront, a collaborative research group dedicated to the study of non-Alzheimer disease degenerative dementias, from the National Health and Medical Research Council of Australia program grant (#1037746 and #1095127)..

CO is supported by a National Health and Medical Research Council Neil Hamilton Fairley Fellowship (1091310) and by the Wellcome Trust (200181/Z/15/Z).

